# Cloning and functional expression of a food-grade circular bacteriocin, plantacyclin B21AG, in probiotic *Lactobacillus plantarum* WCFS1

**DOI:** 10.1101/2020.04.23.057653

**Authors:** Mian Chee Gor, Aida Golneshin, Thi Thu Hao Van, Robert J. Moore, Andrew T. Smith

**Affiliations:** School of Science, RMIT University, Bundoora, Victoria, Australia; Edlyn Foods Pty Ltd, Melbourne, Victoria, Australia; Griffith Sciences, Griffith University, Southport, Queensland, Australia

## Abstract

There is an increasing consumer demand for minimally processed, preservative free and microbiologically safe food. These factors, combined with risks of antibiotic resistance, have led to interest in bacteriocins produced by lactic acid bacteria (LAB) as natural food preservatives and as protein therapeutics. We previously reported the discovery of plantacyclin B21AG, a novel circular bacteriocin produced by *Lactobacillus plantarum* B21. Here, we describe the cloning and functional expression of the bacteriocin gene cluster in the probiotic *Lactobacillus plantarum* WCFS1. Genome sequencing demonstrated that the bacteriocin is encoded on a 20 kb native plasmid, designated as pB21AG01. Seven open reading frames (ORFs) putatively involved in bacteriocin production, secretion and immunity were cloned into an *E. coli*/*Lactobacillus* shuttle vector, pTRKH2. The resulting plasmid, pCycB21, was transformed into *L. plantarum* WCFS1. The cell free supernatants (CFS) of both B21 and WCFS1 (pCycB21) showed an antimicrobial activity of 800 AU/mL when tested against the WCFS1 (pTRKH2) indicator strain, indicating functional expression of plantacyclin B21AG. Real-time PCR analysis revealed that the relative copy number of pB21AG01 was 7.60 ± 0.79 in *L. plantarum* B21 whilst pCycB21 and pTRKH2 was 0.51 ± 0.05 and 25.19 ± 2.68 copies, respectively in WCFS1. This indicates that the bacteriocin gene cluster is located on a highly stable, low copy number plasmid pB21AG01 in *L. plantarum* B21. Inclusion of the native promoter for the bacteriocin operon from pB21AG01 may result in similar inhibitory zones observed in both wild type and recombinant hosts despite the low copy number of pCycB21.

## Introduction

Bacteriocins are ribosomally synthesised, extracellularly released peptides or peptide complexes that possess antibacterial activity against species usually closely related to the producer strains or a wider range of microorganisms [1,2]. Interestingly, the bacteriocins produced by gram-positive bacteria seem to exhibit broader spectrum activity compared to the gram-negative bacteria [3]. Among the gram-positive bacteria, bacteriocins produced by the food-grade lactic acid bacteria (LAB) have attracted considerable interest because they are generally regarded as safe (GRAS). Being proteins they can be easily degraded by proteases in the mammalian gastrointestinal tract, making them safe for human consumption and minimizing the risk of developing resistant bacteria [4,5]. They have been widely used as natural food preservatives for controlling food-borne and food-spoilage bacteria without affecting sensory qualities. They also have huge potential in veterinary applications and as next-generation antibiotics against multi-drug resistant (MDR) pathogens [5–7]. One of the advantages of bacteriocins over conventional antibiotics is that they are directly gene encoded, making bioengineering feasible to enhance their productivity or specificity towards target pathogens [5,8].

The classification of bacteriocins produced by gram-positive bacteria has been constantly revised due to the extensive research performed over the last two decade [9–11]. Here we use the classification proposed by Acedo et al. [12]. Class I contains modified peptides including lantibiotics, lipolanthines, linear azol(in)e-containing bacteriocins, thiopeptides, bottromycins, sactibiotics, lasso peptides, glycocins and circular bacteriocins. Class II are unmodified peptides such as YGNG-motif containing bacteriocins, two-peptide bacteriocins, leaderless bacteriocins and other linear bacteriocins. Class III are large heat labile bacteriocins such as bacteriolysins, non-lytic large bacteriocins and tailocins. Of these, the circular bacteriocins have gained considerable attention as they generally exhibit broad antimicrobial activity. They are synthesised as linear pre-peptides where the leader peptides of variable sizes (3 – 35 amino acids) are cleaved off during maturation, forming 58 – 70 amino acid peptides which are covalently linked by a largely unknown cyclisation mechanism [13,14]. The circular structures appear to enhance their pH and thermal stability as well as protease resistance. These properties make them a preferred candidate for potential industrial applications compared to the other classes of bacteriocins [13,15].

Among the LAB, bacteriocins produced by *Lactobacillus*, in particular *Lactobacillus plantarum* have been widely studied for several reasons. *L. plantarum* is a versatile species that is widely found in a variety of sources, including meat, dairy, fish, fruit and vegetables [16]. It is also one of the natural inhabitants of the human gastrointestinal tract (GIT) where its ability to survive passage through the GIT makes it an attractive vector for vaccine delivery [17,18]. The availability of the complete genome sequence of *L. plantarum* WCFS1 and genome mining tools have facilitated the characterisation of the genetic organisation of the plantacyclin (*pln*) loci from this species [19]. Hitherto, several other class II linear two-peptide bacteriocins produced by *L. plantarum* strains have been described. For example, plantaricin C-19 produced by *L. plantarum* C-19, isolated from fermented cucumber, and plantaricin NA produced by *L. plantarum,* isolated from ‘ugba’, an African fermented oil-bean seed showed strong antimicrobial activity against the food-borne pathogen, *Listeria monocytogenes* [20,21]. Bacteriocin AMA-K produced by *L. plantarum* AMA-K, isolated from fermented milk exhibited strong adsorption to cells of *L. monocytogenes*, *L. ivanovii* subsp. *ivanovii* and *L. innocua* [22]. Plantaricin ST8KF produced by *L. plantarum* ST8KF, isolated from kefir, demonstrated antimicrobial activity against *L. casei*, *L. salivarius*, *L. curvatus*, *Enterococcus mundtii* and *L. innocua* [23,24]. In contrast, only one circular bacteriocin, plantaricyclin A produced by *L. plantarum* NI326 has been reported to date. Similarly, this circular bacteriocin is active against beverage-spoilage bacterium *Alicyclobacillus acidoterrestris* [25]. These antimicrobial peptides appear to have great potential in food preservation, particularly in controlling food-borne pathogens. Discovery of more circular bacteriocins is highly favourable over linear peptides due to their superior stability against various stresses [26].

In recent years, research on bacteriocins has progressed from producing the inhibitory compounds in native systems to heterologous production in diverse producer organisms which have the potential to be employed as starters, protectors and/or probiotics [27]. Several strategies for heterologous expression of bacteriocins have been investigated either for overproduction of the bacteriocin or structure-function studies [7,27–29]. We previously reported the discovery of plantacyclin B21AG, a food-grade circular bacteriocin produced by *Lactobacillus plantarum* B21 [30,31]. It is shown to be active against food-borne pathogens including *Clostridium perfringens* and *Listeria monocytogenes*; food spoilage bacteria such as *L. arabinosus*; as well as other LAB including *L. plantarum*, *L. brevis* and *Lactococcus lactis* [30–32]. This study aimed to transfer the production of the broad antimicrobial spectrum of plantacyclin B21AG to a probiotic strain, *L. plantarum* WCFS1 [33]. We demonstrated that the bacteriocin gene cluster can be recombinantly expressed in *L. plantarum* WCFS1 at a level comparable to the native producer *L. plantarum* B21. The mobilization of the plantacyclin B21AG operon into the probiotic, *L. plantarum* WCFS1, enhances the antimicrobial activity spectrum of the strain, potentially making it more useful for use in the food industry and for clinical applications.

## Materials and methods

### Bacterial strains, plasmids and culture conditions

Bacterial strains and plasmids used in this study are listed in Table 1. All *Lactobacillus* strains were cultured statically in deMan, Rogosa and Sharpe (MRS) broth (Becton, Dickinson and Company, USA) at 37 °C under aerobic conditions. *Escherichia coli* strains were grown in Luria Bertani (LB) medium (Becton, Dickinson and Company, USA) at 37 °C with continuous agitation at 250 rpm. For selection, medium were supplemented with 100 μg/mL of ampicillin and/or 150 μg/mL of erythromycin for *E. coli* and 15 μg/mL of erythromycin for *Lactobacillus*.

**Table 1.**
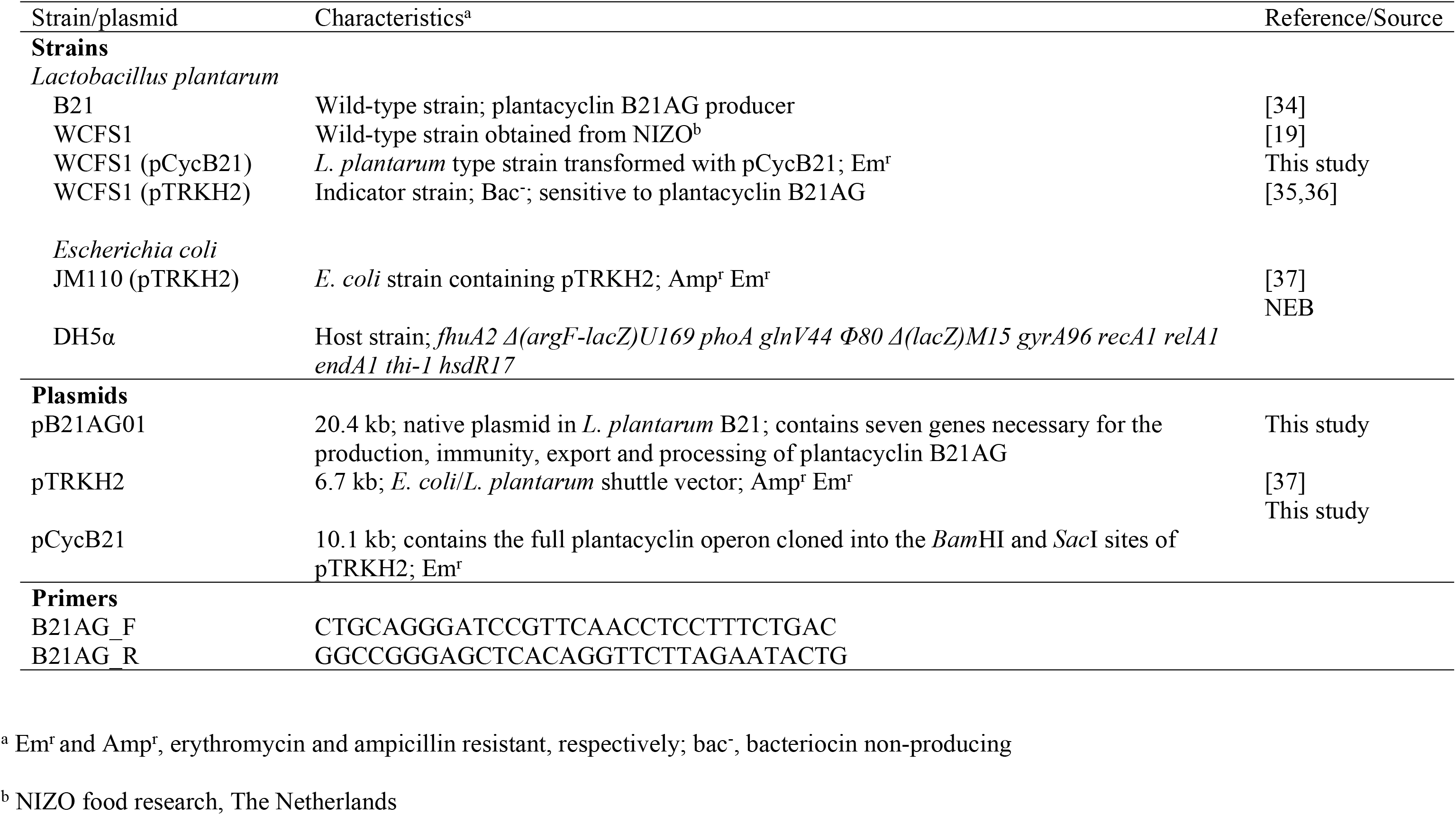
Bacterial strains, plasmids and primers.

**Table 2.**
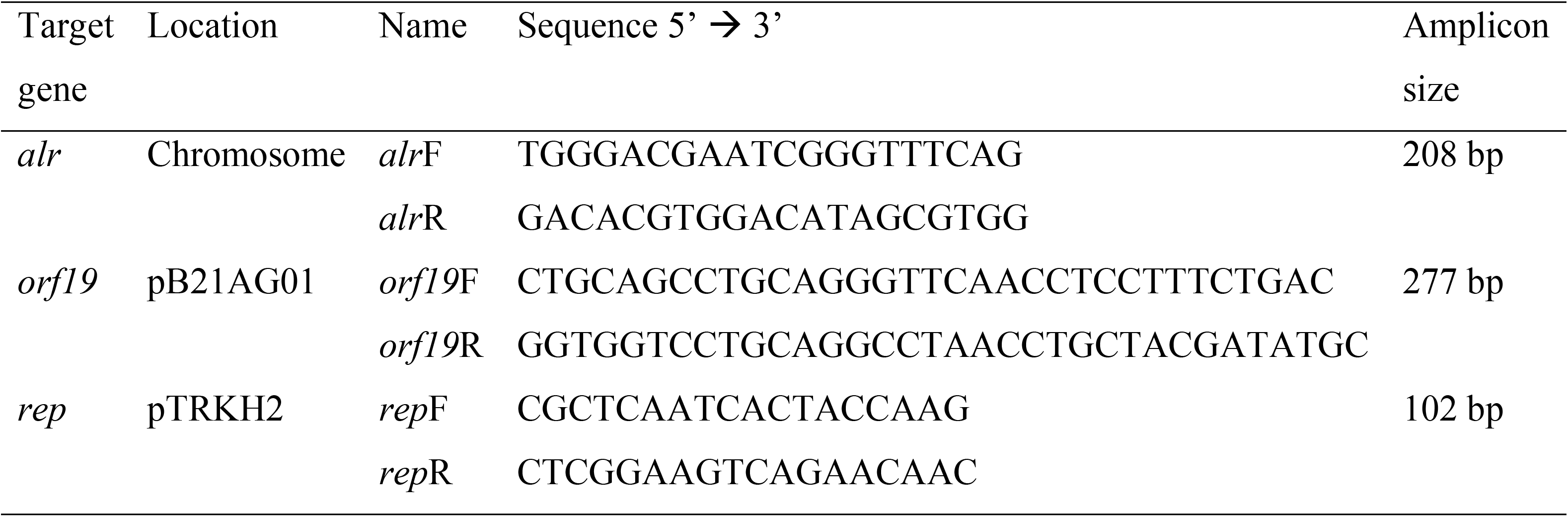
Primers used for plasmid copy number (PCN) detection with real-time PCR.

### Sequence determination and genetic analysis of pB21AG01

The full genome sequence of *L. plantarum* B21, including the 20.4 kb plasmid, pB21AG01, was sequenced at the Beijing Genomics Institute (BGI) using the Illumina HiSeq 2000 platform (Illumina, USA) and assembled with SOAPdenovo software [38]. The plasmid was annotated using RAST [39]. The resulting open reading frames (ORFs) were confirmed using blastp against the NCBI non-redundant protein database [40]. The obtained putative protein sequences were searched for conserved domains using the NCBI Conserved Domain Database (CDD) site [41] and also examined for transmembrane domains using the TMHMM transmembrane prediction algorithm at http://www.cbs.dtu.dk [42].

### DNA manipulations, plasmid constructions and E. coli transformations

Total gDNA from *E. coli* and *Lactobacillus* was isolated using the GeneElute™ Bacterial Genomic DNA Kit (Sigma-Aldrich, USA) as described in the user manual. Plasmids from *E. coli* were extracted using the ISOLATE II Plasmid Mini kit (Bioline, Australia) according to manufacturer’s instruction. Plasmids from *Lactobacillus* were prepared using QIAGEN^®^ Plasmid Midi Kit (Qiagen, Germany) following supplier’s direction with a few modifications to cell wall lysis. One hundred millilitre of overnight culture was harvested by centrifugation and washed in 20 mL STE buffer (6.7 % sucrose; 50 mM Tric-HCl, pH 8.0; 1 mM EDTA) [43] to remove and neutralise acids produced during cell growth. The bacterial pellet was then resuspended in 4 mL STE buffer containing 10 mg/mL lysozyme and incubated at 37 °C for 1 hour.

For the construction of pCycB21, the full plantacyclin B21AG operon with its native promoter was amplified from *L. plantarum* B21 plasmids using One*Taq*^®^ 2X Master Mix with Standard Buffer (NEB, USA) in a T100™ Thermal Cycler (Bio-Rad, USA). PCR, using the primers indicated in Table 1, was performed as follows: initial denaturation for 30 s at 94 °C; followed by 30 cycles of denaturation for 30 s at 94 °C, annealing for 30 s at 55 °C and extension for 4 min at 68 °C; and a final extension for 10 min at 68 °C. The amplified product was purified using the ISOLATE II PCR and Gel Kit (Bioline, Australia) and cloned into the *Bam*HI and *Sac*I sites of the *E. coli*/*Lactobacillus* shuttle vector, pTRKH2. The construct was transformed into *E. coli* DH5α (NEB, USA) according to manufacturer’s protocol in order to obtain sufficient amount of plasmid DNA for subsequent transformation into LAB. The recombinant plasmid is confirmed by PCR, double restriction enzyme digestion and DNA sequencing.

### Electroporation of LAB

Electroporation of LAB was performed as described by Mason et al. [44] with a few modifications. Briefly, 8 mL of overnight LAB cultures were diluted into 40 mL of fresh pre-warmed MRS broth containing 2 % glycine. The diluted culture was incubated for 1.5 hr at 37 °C. The cells were pelleted by centrifugation at 4000 x g for 2 min at 4 °C and washed with 40 mL of ice-cold Milli-Q water. Cells were then resuspended in 40 mL of ice-cold 50 mM EDTA and incubated on ice for 5 min. Centrifugation was repeated followed by washing the cells in 40 mL of ice-cold 0.3 M sucrose. The cells were resuspended in 200 μL of 0.6 M sucrose. Finally, 3 μg of DNA in 50 μL of sterile Milli-Q water was added to 50 μL of freshly prepared competent cells and transferred into a pre-chilled electrocuvette with a 0.2-cm electrode gap (Cell Projects, UK). The cell suspension containing plasmid DNA was electroporated using a Gene Pulser electroporator (Bio-Rad, USA) with the following parameters: 1.5 kV, 200 Ω parallel resistance and 25 μF capacitance. The cells were transferred immediately after electroporation into 1.3 mL of pre-warmed MRS broth and incubated for 3 hrs at 37 °C. Two hundred microliters of the cells were plated onto MRS agar containing erythromycin and incubated for 2 days at 37 °C. Recombinant plasmids were confirmed by PCR and double restriction enzyme digestion.

### Antimicrobial activity assay

The antimicrobial activity of the bacteriocin produced by LAB was evaluated using the well diffusion agar (WDA) method [45]. Briefly, cell free supernatants (CFS) of *L. plantarum* B21 and WCFS1 (pCycB21) were harvested from 15 mL overnight LAB culture by centrifugation at 4,000 x g for 20 min at 4 °C. The CFS was then concentrated 15-fold using an Amicon^®^ Ultra-15 Centrifugal Filter Devices (Merck Millipore, Germany) and stored at 4 °C until used. To evaluate the antimicrobial activity of plantacyclin B21AG, MRS agar plates supplemented with 10 μg/mL of erythromycin were seeded with 10^6^ cfu/mL of *L. plantarum* WCFS1 (pTRKH2), used as the indicator strain. Wells were made in the agar using a sterile 8-mm cork borer. One hundred microliters of the 2-fold serial diluted CFS was then loaded into the wells and the plates were incubated at 30 °C for 16 – 18 h. Antimicrobial activity was expressed as arbitrary unit (AU/mL) using the following equation, a^b^ x 100, where “a” is the dilution factor, “b” is the last dilution showing an inhibition zone of at least 2 mm in diameter [46].

### Extraction of Plantacyclin B21AG with 1-butanol

Plantacyclin B21AG was purified from *L. plantarum* B21 and *L. plantarum* WCFS1 (pCycB21) using 1-butanol as described by Abo-Amer [47] with the following modifications: the concentrated CFS was mixed with ½ volume of water-saturated butanol for 20 s. The mixture was incubated at room temperature for 10 min to allow phase separation before centrifugation at 10,000 x g for 10 min. The butanol phase was transferred to a clean 1.5 mL tube whilst the aqueous phase was subjected an additional butanol extraction. The two butanol fractions containing plantacyclin B21AG were combined and the solvent was removed using a freeze dryer (FDU-8612, Operon Co. Ltd, Korea). The lyophilised protein was dissolved in 20 mM sodium phosphate buffer (pH 6.0).

### Mass spectrometry analysis

The protein was subjected to matrix-assisted laser desorption/ionization time-of-flight mass spectrometry (MALDI-TOF-MS) analyses as described by Vater et al. [48]. The MALDI-TOF mass spectra were recorded using an Autoflex Speed MALDI-TOF instrument (Bruker, Germany) containing a 355 nm Smartbeam II laser for desorption and ionization. 10 mg of α-cyano-4-hydroxycinnamic acid dissolved in 70 % acetonitrile (ACN) containing 0.1 % (v/v) trifluoroacetic acid (TFA) was used as matrix solution. Five microliters of bacteriocin samples were mixed with equal volume of matrix solution and 1 μL of the mixture was spotted onto the target, air dried and measured.

### Plasmid copy number determination by real-time PCR

The copy number of the native (pB21AG01) and recombinant (pCycB21) plasmids were determined using real-time PCR according to Škulj et al. [49]. A 5-fold serial dilution of total DNA extracted from *L. plantarum* B21 was used for the standard curves (final 1 ng/μL to 0.0016 ng/μL). Real-time PCR reactions were performed in 12 μL mixtures containing 1 x SensiFAST SYBR No-ROX mix (Bioline, Australia), 400 nM of each forward and reverse primer (Table 3) and 1 μL of DNA. The alanine racemase gene (*alr*), a single copy, chromosomal gene from *L. plantarum* WCFS1, was selected as the reference gene (GeneBank Accession No. AL935253) whilst the bacteriocin structural (*orf19*) was chosen as the target for detection of the recombinant plasmid pCycB21. The replication (*rep*) gene was used as the target to detect pTRKH2 in WCFS1. Separate reactions were prepared for the detection of chromosomal and plasmid amplicons. All reactions were performed in duplicate using the Rotor-Gene™ Q (Qiagen, Germany). Thermocycling conditions were: initial denaturation for 3 min at 95 °C, followed by 40 cycles of 5 s at 95 °C, 10 s at 55 °C and 20 s at 72 °C. Fluorescence signal was acquired at the end of each 72 °C step.

**Table 3.**
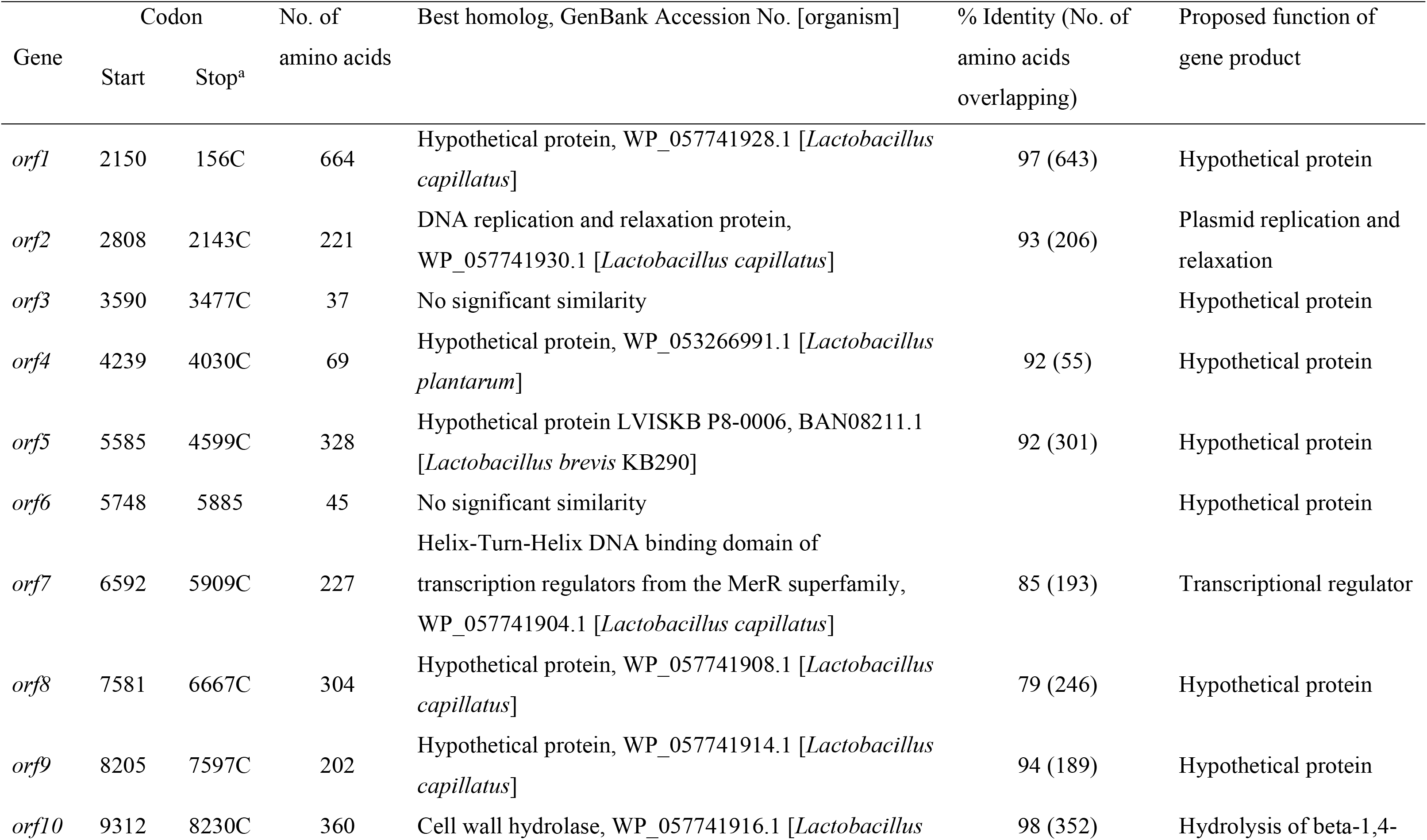
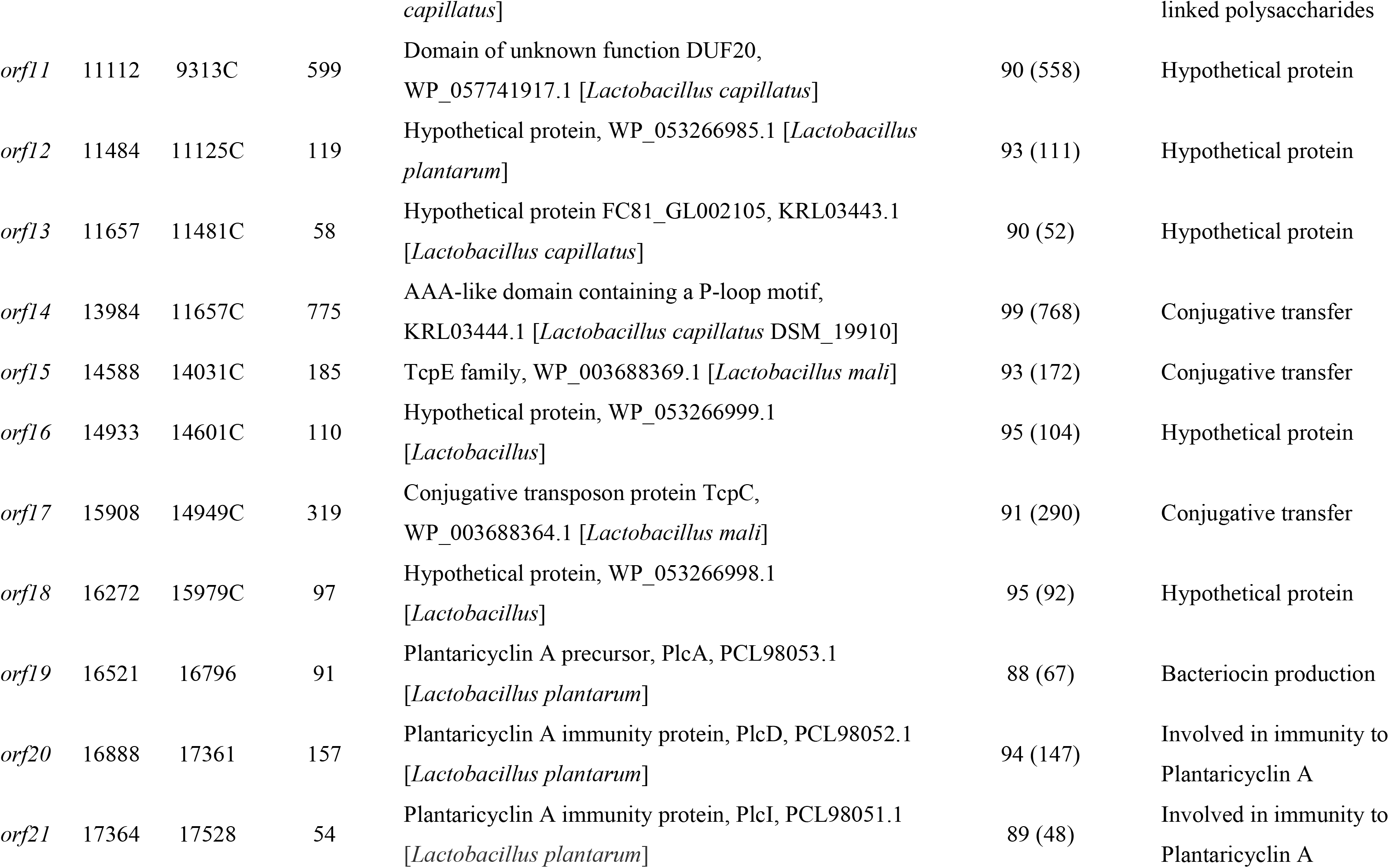
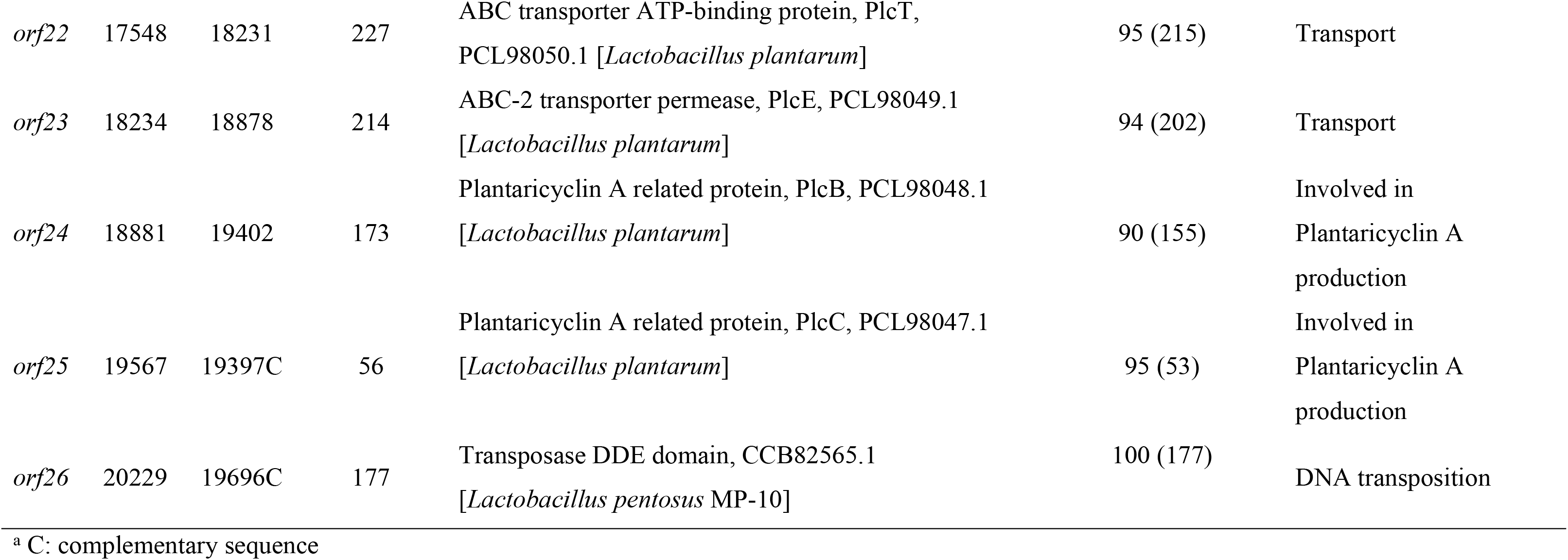
Putative genes and their proposed function deduced from the amino acid sequences of pB21AG.

The slope of the relative standard curve with a condition that r^2^ > 0.99 was used to calculate the amplification efficiency (E) using equation 1.

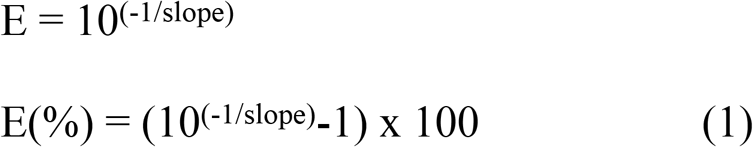

The plasmid copy number (PCN) was calculated based on equation 2 using efficiency (E) and Ct values for both chromosomal (c) and plasmid (p) amplicons.

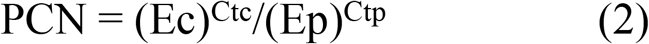

## Results

### Sequence analysis of pB21AG01

Sequence analysis revealed that *L. plantarum* B21 (GenBank Accession No. CP010528) harboured at least two cryptic plasmids, designated as pB21AG01 (GenBank Accession No. CP025732) and pB21AG02 (GenBank Accession No. CP025733). For the purpose of this study, we focused our analysis on pB21AG01 as it was found to encode the genes responsible for the production of a circular bacteriocin. Plasmid profile analysis revealed that pB21AG01 is a 20,429 bp circular DNA molecule with a GC content of 37.3 %. A total of 26 open reading frames (ORF) were identified (Table 3). 14 ORFs of the pB21AG01 were homologous to proteins with known or predicted functions whilst the remaining 12 ORFs were either homologous to hypothetical proteins lacking functional predicts or had no significant homology with any protein sequences in the GenBank databases.

Seven ORFs were predicted to encode genes putatively responsible for the production, immunity and transport of plantacyclin B21AG (*orf19* – *orf25*) [30]. *Orf19* showed 88 % identity to Plantaricyclin A precursor (PlcA), a circular bacteriocin produced by *L. plantarum* (PCL98053.1), presumably the structural gene responsible for the production of plantacyclin B21AG. It encodes 91 amino acids consisting of a 33 amino acid leader peptide and a 58 amino acid bacteriocin mature peptide, which is 86 % identical to *plcA*, the structural gene of Plantaricyclin A [25]. *Orf20* and *orf21* are 94 % and 89 % identical to *plcD* (PCL98052.1) and *plcI* (PCL98051.1), putatively involved in immunity to plantaricyclin A. *Orf22* and *orf23* are possible transporters of the plantacyclin B21AG. *Orf22* showed 95 % identity to *plcT*, a gene encoded for ABC transporter ATP-binding protein in *L. plantarum* (PCL98050.1) whilst *orf23* is 94 % identical to *plcE*, an ABC-2 transporter permease encoding gene in *L. plantarum* (PCL98049.1), respectively. *Orf24* and *orf25* are predicted as plantaricyclin A-related proteins, putatively involved in the production of the circular bacteriocin. Amino acid sequence comparison with gassericin A revealed that *orf20*, *orf24* and *orf25* may be membrane associated proteins, but their roles in bacteriocin production/immunity remain unknown [50]. Transmembrane analysis using TMHMM revealed the presence of 2 transmembrane domains in *orf19*, *orf21* and *orf25*. *Orf20* and *orf23* and *orf24* contain 4, 6 and 5 transmembrane domains, respectively. This result is consistent with the properties of the proteins involved in gassericin A production. No transmembrane domain was predicted for *orf22*.

Although no *rep* gene or direct repeats were found in pB21AG01, *orf2* showed 93 % identity to a protein in *L. capillatus* (WP_057741930.1) which is essential for plasmid replication and relaxation. In addition, *orf7* is predicted to be a Helix-Turn-Helix DNA binding domain of transcription regulators from the MerR superfamily, accession number WP_057741904.1 [*Lactobacillus capillatus*] [41]. They have been shown to mediate responses to environmental stress including exposure to heavy metals, oxygen radicals and antibiotics or drugs in a wide range of bacterial genera [51]. *Orf10* is predicted as cell wall hydrolase, accession number WP_057741916.1 [*Lactobacillus capillatus*]. It is putatively involved in the hydrolysis of beta-1,4-linked polysaccharides [41]. *Orf26* is 100 % identical to the pfam01609 transposase DDE domain of *Lactobacillus pentosus* MP-10 (CCB82565.1) which is essential for efficient DNA transposition. Although we did not find any *tra* genes responsible for bacterial conjugation in pB21AG01, three ORFs were annotated as genes related to plasmid mobilisation. *Orf15* and *orf17* showed 93 % and 91 % identity to genes encoded for the conjugative transposon proteins TcpE (WP_003688369.1) and TcpC (WP_003688364.1) of *L. mali*, respectively. *Orf14* is 99 % identical to the pfam 12846 AAA-like domain containing a P-loop motif from *L. capillatus* DSM_19910 (KRL03444.1), putatively involved in conjugative transfer [41].

### Cloning of the B21AG gene cluster and LAB transformation

The 3,424 bp sequence corresponding to seven genes putatively involved in plantacyclin B21AG production, secretion and immunity was cloned into the pTRKH2 shuttle vector. The resulting plasmid, pCycB21, transformed into *L. plantarum* WCFS1 is detailed in Fig 1.

**Fig. 1.**
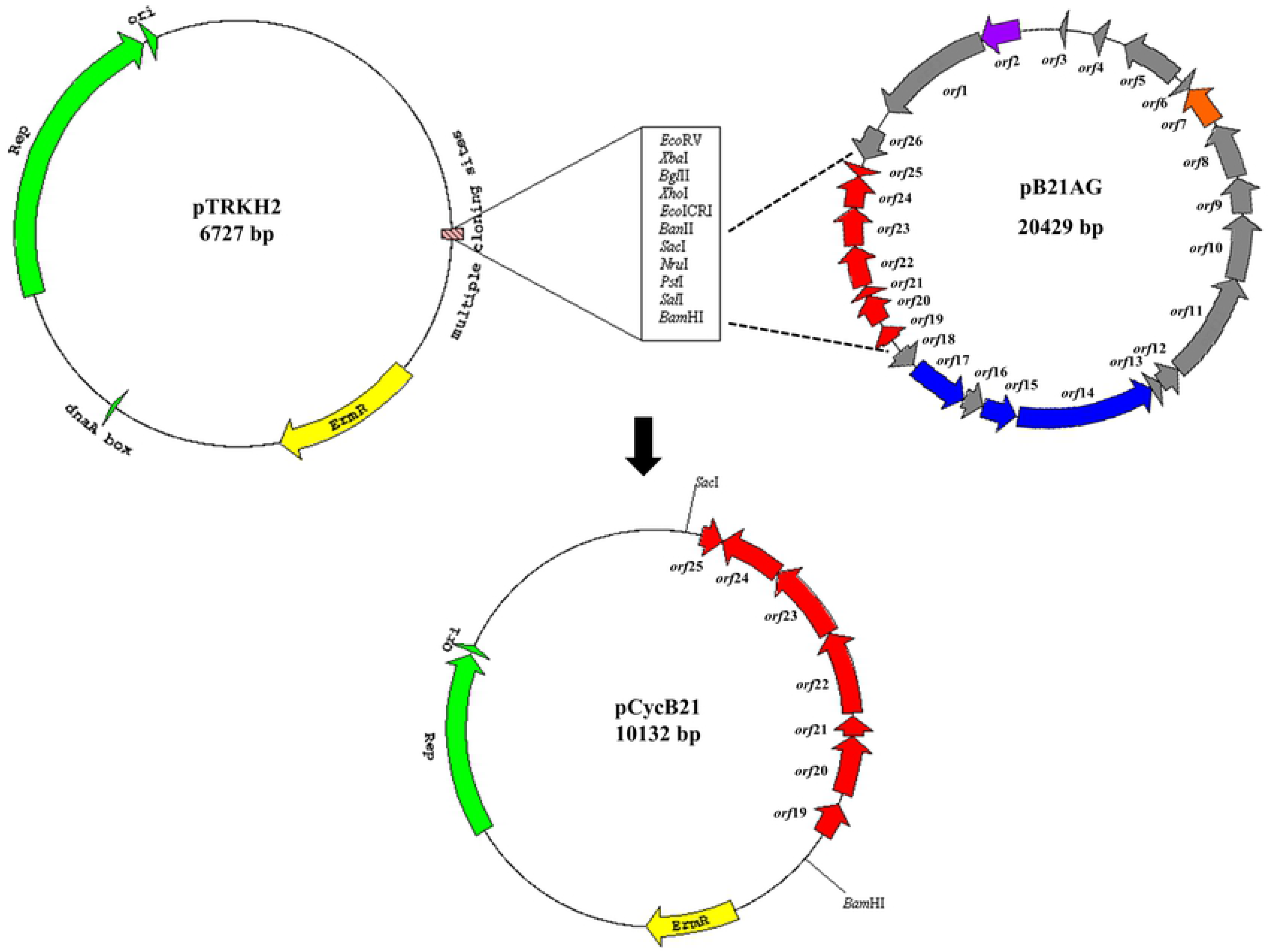
Map of pTRKH2, pB21AG and pCycB21. A: Map of pTRKH2 consisting an origin of replication from plasmid P15A and a replication gene (green arrows) for replication in both *E. coli* and gram positive bacteria; an erythromycin resistance gene (ErmR, yellow arrow) for selection in *E. coli* and LAB; and a multiple cloning site (MCS). B: Map of pB21AG containing 26 open reading frames, with seven ORFs corresponding to bacteriocin production, immunity and transportation (red arrows); a plasmid replication and relaxation gene (purple arrow); a transcription regulator (orange arrow), three conjugation transfer genes (blue arrows) and 14 other ORFs (grey arrows). C: Map of pCycB21 harbouring seven bacteriocin-associated genes cloned into *Sac*I and *Bam*HI restriction sites.

The pTRKH2 and pCycB21plasmids were electrotransformed into *L. plantarum* WCFS1 with efficiencies of 2.4 × 10^2^ and 3.4 × 10^2^ transformants per μg DNA, respectively. Addition of glycine in the growth medium inhibits formation of cross-linkages in the cell wall where _L_-alanine is replaced by glycine, thereby weakening the cell wall and facilitating DNA update by the cells [52,53].

### Assay of plantacyclin B21AG expression

Antimicrobial activities of the wild type *L. plantarum* B21 and recombinant host *L. plantarum* WCFS1 were assayed using the well diffusion agar (WDA) method. Since pTRKH2 conferred erythromycin (Em) resistance on WCFS1, culture supernatants containing Em were assayed using indicator WCFS1 (pTRKH2) without removing the antibiotics. To eliminate the effect of acid production, the pH of cell free supernatants were neutralised to pH6.5. Both wild type and recombinant hosts were found to produce inhibition zones against the indicator strain up to 1:8 dilution (Fig 2). The antimicrobial activity was calculated as approximately 800 AU/mL for both strains. This indicates that the inhibitory activity is not due to acid production but to an antimicrobial substance secreted into the broth [54]. The CFS of control cultures containing WCFS1 (pTRKH2) did not show any inhibitory activity, confirming that the recombinant plasmid pCycB21 was responsible for the antimicrobial activity.

**Fig. 2.**
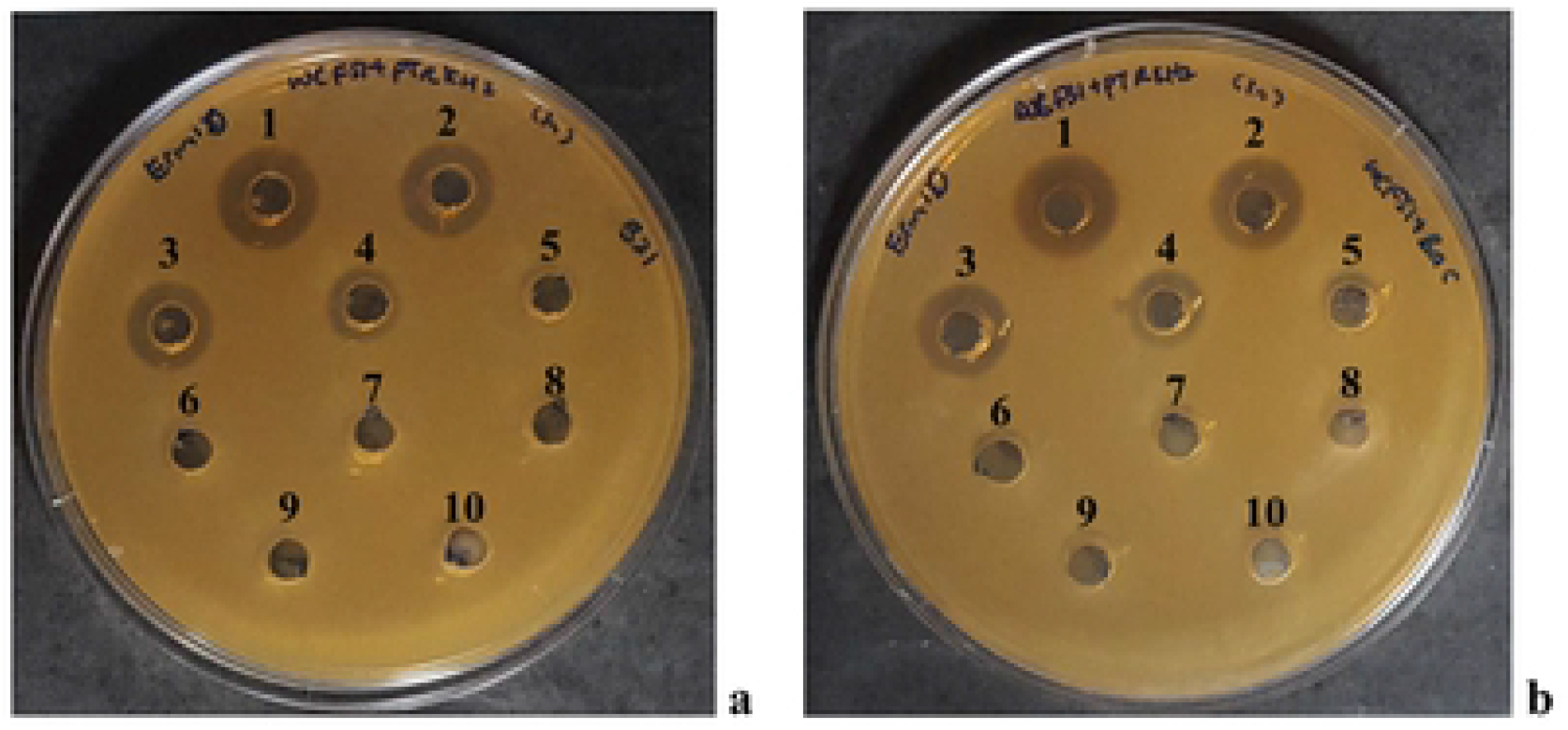
Antimicrobial activity of a two-fold serial dilution of Plantacyclin B21AG secreted by (a) *Lactobacillus plantarum* B21 and (b) WCFS1 harbouring pCycB21. *L. plantarum* WCFS1 (pTRKH2) was used as the indicator strain. Numbers above the wells correspond to the CFS dilution in each well. 1, Undiluted CFS; 2, 1:2 dilution of CFS; 3, 1:4 dilution of CFS; 4, 1:8 dilution of CFS; 5, 1:16 dilution of CFS; 6, 1:32 dilution of CFS; 7,1:64 dilution of CFS; 8, 1: 128 dilution of CFS. Well 9 (indicator strain) and well 10 (MRS broth) are negative control.

### Mass spectrometry analysis

The plantacyclin B21AG produced by the wild type B21 and recombinant host WCFS1 (pCycB21) was purified by extraction into butanol. MALDI-TOF-MS analysis revealed a major peptide of molecular mass of 5663.9 Da, essentially identical to the plantacyclin B21AG produced by the wild type *L. plantarum* B21 (5664.7 Da) (Fig 3). No major peaks were observed for WCFS1 transformed with the shuttle vector pTRKH2, corroborating the results from the functional expression assay described previously (Fig 2, well 9).

**Fig. 3.**
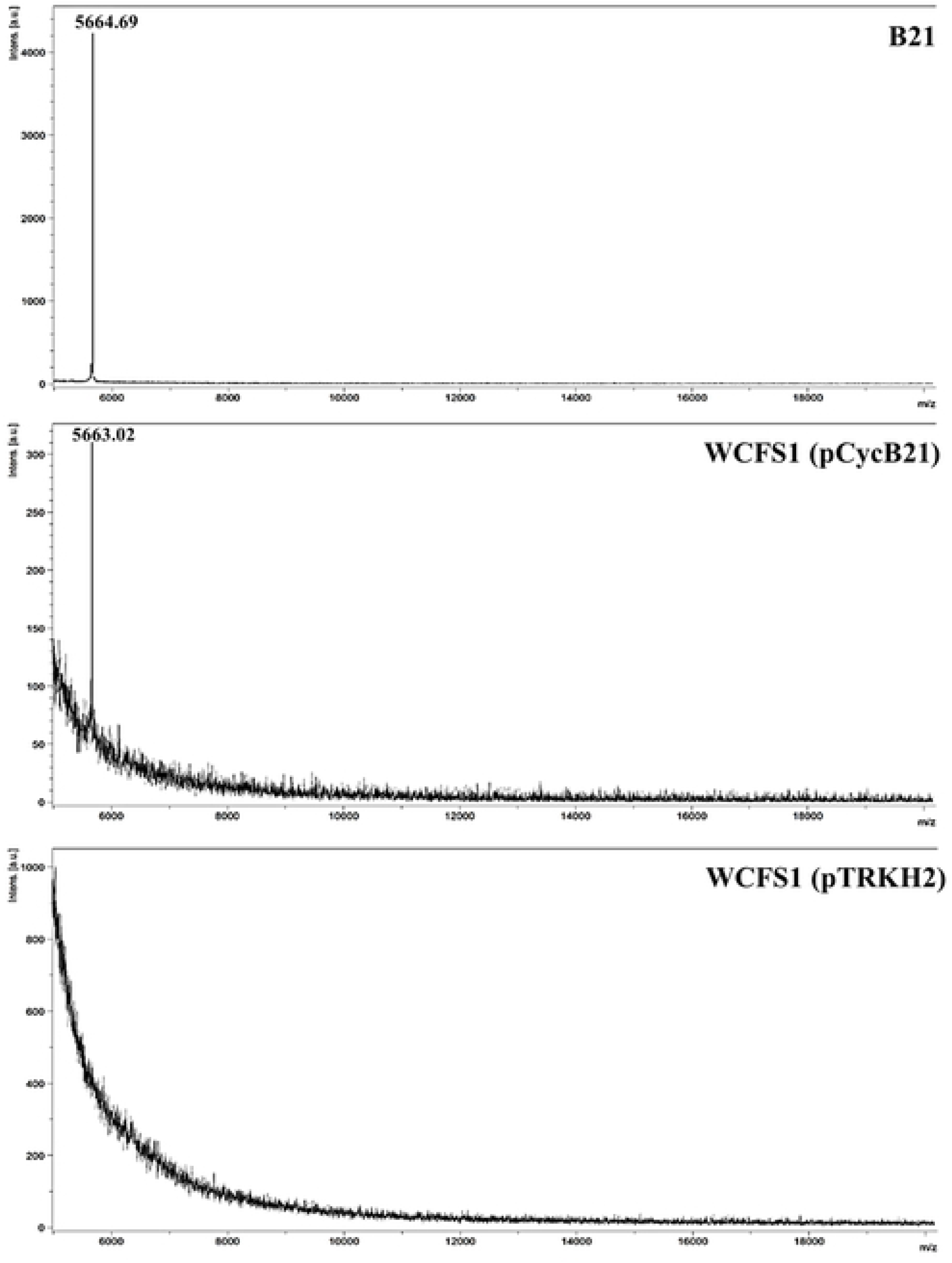
MALDI-TOF-MS spectrum of plantacyclin B21AG, showing a single peak at molecular mass of 5664.69 for B21 and 5663.92 for WCFS1 (pCycB21). No major peaks were observed for WCFS1 (pTRKH2).

### Copy number of pB21AG01 and pCycB21

The relative copy number of pB21AG01 and pCycB21 was determined by real-time PCR using the single copy alanine racemase (*alr*) as reference gene. In our experiment, the standard curves obtained for *alr*, *orf19* and *rep* were linear (R^2^ > 0.99) over the range tested; whilst the amplification efficiency for all experiments ranged between 90 – 102 %, which is within the acceptable range (90-110%) [55]. Analysis of the results revealed that approximately 7.6 ± 0.79 copies per cell of pB21AG01 was detected in *L. plantarum* B21 whilst pCycB21 in *L. plantarum* WCFS1 was present at a noticeably lower level of just 0.5 ± 0.05 copies, indicating that only half the cells carry the recombinant plasmid. In contrast, the copy number of the shuttle vector pTRKH2 in *L. plantarum* WCFS1 was high at 25.19 ± 2.68 copies per cell (average for two clones ± standard deviation) per chromosome equivalent.

### Discussion

Several lactic acid bacteria species have been recognised as probiotics that possess important traits such as the production of bacteriocins and organic acids, adhesion to host cells, and resistance to antibiotics and heavy metals [56,57]. A number of native plasmids that encode these probiotic traits have been sequenced from *L. plantarum* [56,58], *L. salivarius* [59] and *L. fermentum* [57]. In this study, seven bacteriocin-associated genes were found to be located on a 20 kb native plasmid, pB21AG01 in *L. plantarum* B21. No replication protein (repB) or initiator replication family protein (repA) was found in pB21AG01. However, a DNA replication and relaxation conserved domain was detected in *orf2*. We could not detect any clear repeats in the region upstream of *orf2*, suggesting that the plasmid may replicate through a mechanism yet to be determined [60].

In addition, we identified two *tcp* loci, TcpE (*orf15*) and TcpC (*orf17*), which are involved in the transfer of conjugative plasdmid, pCW3 from *Clostridium perfringens*. TcpE was shown to play a role in the formation of Tcp transfer apparatus in the gram positive *Clostridium perfringens* [61]. TcpC was identified as a bitopic membrane protein, where membrane localisation is important for its function, oligomerisation and interaction with other conjugation proteins [62]. Bantwal et al. [63] proposed that TcpC may initiate accumulation of peptidoglycan hydrolase at the cell wall of *Clostridium*, resulting in degradation of peptidoglycan, thus facilitate the formation of transfer apparatus. Interestingly, a cell wall hydrolase (*orf10*) was also found in pB21AG01, suggesting that TcpC may play a role in promoting the hydrolysis of *Lactobacillus* cell wall and subsequently the transfer of pB21AG01. However, we did not transfer the plasmid by mating because we could not determine a full set of genes responsible for the mobilisation of the plasmid. pCW3 has a novel conjugation region consisting of 11 genes encoding the Tcp proteins (TcpA, TcpB, TcpC, TcpD, TcpE, TcpF, TcpG, TcpH, TcpI, TcpJ and TcpM) [61,64]. Several studies have demonstrated that TcpA, TcpD, TcpE, TcpF and TcpH are essential to form the conjugation complex. Moreover, we could not detect any antibiotic resistance and/or heavy metal resistance genes in pB21AG01, which could be used as natural selection markers if we were to transfer the plasmid by mating. Due to the lack of *tcp*-encoded proteins and selection markers, we decided to transfer the plasmid by electroporation.

Electroporation seems to be an efficient method to transfer plasmid DNA into LAB to enhance their probiotic functionality, or to secrete therapeutic proteins into the culture medium for human and animal health [65]. However, the success rate of LAB transformation is extremely low compared to *E. coli* due to various restriction modification (RM) system encoded by the host. RM systems are required to protect bacteria from foreign DNA such as genetically transferred plasmid DNA or the bacteriophage DNA [65,66]. Since DNA manipulation is easier in *E. coli* than in *Lactobacillus*, we have built the recombinant plasmid in the shuttle vector pTRKH2, followed by propagating the plasmids in *E. coli* to obtain sufficient amount of plasmid DNA for LAB transformation. Numerous attempts at electrotransformation were performed according to various protocols described in the literature but without any success. The parameters for electrotransformation that we have tried included varying the percentage of glycine added to LAB culture prior to pelleting the cells; addition of PEG_1500_ and EDTA in the washing steps; various concentrations of plasmid DNA; different electroporation buffers and diverse combinations of voltage (V), capacitance (μF) and resistance (ohm). The final modified version of the method described [44], that we eventually had success with is described in the materials and methods section.

A few attempts have been made to heterologously expressed bacteriocins in different LAB species because they promise a food grade background, where the expression of bacteriocins would enhance their probiotic functionality [28,67]. However, all of these peptides that have been successfully expressed heterologously belong to the class II bacteriocins. To date, only one circular bacteriocin, plantaricyclin A from *L. plantarum* NI326, has been successfully cloned into a nisin-inducible plasmid and expressed in *L. lactis* pNZPlc. Both the recombinant host and the wild type producer exhibited similar level of antimicrobial activity [25], indicating that circular bacteriocin can be heterologously expressed in other LAB species. Our results are in accordance to Borrero et al. [25], where both native producer and the recombinant host expressed similar antimicrobial activity up to 800 AU/mL. This result is also confirmed by a single peak observed in our mass spectrometry analyses. In contrast, *L. plantarum* WCFS1 transformed with the empty vector pTRKH2 produced no bacteriocin activity. These results indicate that the bacteriocin activities observed are due to the cloned genes (*orf19 – orf25*). We demonstrated that a 3.4 kb plasmid region of *L. plantarum* B21 is sufficient for functional expression of plantacyclin B21AG. However, our attempt to transform pCycB21 into *Lactobacillus agilis* La3, a type of LAB found to be colonising chicken gastrointestinal tract (GIT) [68], did not result in heterologous expression of the bacteriocin. This result suggests that the expression of recombinant protein in LAB is species-specific. One possible explanation is that the native promoter cloned is specific to *L. plantarum*, and a promoter from *L. agilis* is probably required for heterologous expression. Although bacteria promoters share similar features, promoter strength is strain- and context-specific, and can vary significantly within LAB [67,69,70].

The isolation and purification of bacteriocins from their LAB producers is often very time-consuming and labour intensive [71]. Many studies have been performed to heterologously express and overexpress the class II bacteriocins in *E. coli* to facilitate the production of these antimicrobial peptides. For instance, sakacin P, pediocin PA-1, divercin V41 and plantaricin NC8 have been successfully expressed in *E. coli* [71–74]. However, no circular bacteriocins have been successfully expressed in *E. coli*. Kawai et al. [75] tried to express a circular bacteriocin, gassericin A, in *E. coli* JM109 as a biotinylated fusion protein. However, a positive clone which accumulated the bacteriocin did not show any antimicrobial activity. Further treatment with factor Xa protease released the N-terminal leader peptide, resulting in an active unclosed gassericin A. The results indicate that expression of circular bacteriocins is host-specific, where a yet-to-be identified host-encoded peptidase is required to cleave the leader peptide, allowing the ligation of N- and C-terminal to happen [13,76].

Plasmid copy number (PCN) analysis showed that the native pB21AG01 is a highly stable, low copy number plasmid in B21. pTRKH2 was selected as a shuttle cloning vector because it has been shown that it is structurally stable in *Lactobacillus vaginalis* Lv5, a common feature of the theta-replicating mechanism [68]. It also has good structural stability in *E. coli*, possibly due to the lack of a resolvase-encoding gene [37]. PCN analysis revealed that pTRKH2 is more stable than pCycB21 in *L. plantarum* WCFS1, indicating that the bacteriocin gene cluster may cause instability of the vector pTRKH2. Erythromycin selection is required to maintain the pCycB21 in *L. plantarum* WCFS1. One possible reason which may contribute to the instability of pCycB21 is plasmid incompatibility, where two plasmids containing the same origin of replication cannot co-exist stably in the cell. Plasmids that have growth advantages, such as faster replication and less toxicity will rapidly outgrow the other plasmids [77]. The host used in this study, *L. plantarum* WCFS1 is known to harbour three native plasmids size 1.9 kb, 2.4 kb and 36 kb [56]. Thus, the introduced bacteriocin gene cluster could be a plausible reason for pCycB21 instability in *L. plantarum* WCFS1. Similarly, the copy number of pCycB21 is extremely low compared to the native plasmid pB21AG01. This suggests that other fitness factors present on the native plasmid pB21AG01 play a role in positive plasmid selection. For instance, apart from the immunity genes which are known to protect bacteriocin-producing strains against its own toxins, the gene encoding for ABC transporter also plays an important role. It has been shown to translocate the bacteriocin across the cytoplasmic membrane, thereby avoiding toxin accumulation in the host cells [78,79]. In our case, the presence of ABC transporter could potentially stabilise pB21AG01 in *L. plantarum* B21. Despite the PCN variation between pB21AG01 and pCycB21, plantacyclin B21AG was expressed at a similar level. The production of plantacyclin B21AG would depend on plasmid stability and copy number differences between pB21AG01 and pCycB21, but more likely, might be caused by the promoters used to drive gene expression [7]. We have cloned the native promoter from pB21AG01 into pCycB21AG, presumably resulting in similar levels of plantacyclin B21AG production. In the future, inducible or controlled promoters may be tested to optimise heterologous production of plantacyclin B21AG [67].

In summary, circular bacteriocins are thought to have more potential to form the next generation of biopreservatives as a consequence of their stability and activity [8]. The ability to transfer vectors harbouring the pB21AG01 gene cluster into an industry standard probiotic *L. plantarum* WCFS1 highlights its biotechnological interest for the overproduction of the antimicrobial peptide with high antimicrobial activity against food-borne pathogens.

## Acknowledgements

This work was supported by RMIT University. We are grateful to Mr. Frank Antolasic for his technical assistance in MALDI-TOF-MS analysis.

